# Validating models of TMS effects with concurrent TMS/fMRI

**DOI:** 10.1101/2023.07.05.547836

**Authors:** Petar I. Petrov, Jord Vink, Stefano Mandija, Nico A.T. van den Berg, Rick M. Dijkhuizen, Sebastiaan F.W. Neggers

## Abstract

The exact dose and spatial pattern of brain activation induced by transcranial magnetic stimulation (TMS) is of importance to all applications of TMS, both clinical and for investigational use. How TMS coils induce electrical fields has been investigated and validated by several groups, but models of how evoked currents interact with neuronal tissue and where activation is induced have hardly been validated empirically.

Here we propose a detailed model of TMS induced currents in the brain building on our previous work on different TMS coil models, taking into account various electromagnetic head tissue properties with finite element modeling and several competing macroscopic models of how brain activation is achieved by the TMS induced electrical field. Importantly, we validate these models using TMS coil discharges administered to the brain of 6 healthy volunteers inside a 3T MRI scanner, while acquiring functional MRI scans at the same time.

We conclude that the mere magnitude of the electric field in the cortical gray matter tissue near the stimulation site provides the best metric to predict neuronal activation as measured by BOLD fMRI, whereas models of brain activation taking into account orientation of E-fields with respect to the cortical columns and layers do not (yet) yield better predictions of neuronal activation induced by TMS.

This report is relevant for neuronavigated TMS approaches attempting to take into account evoked E-fields and neuronal activation to improve precision of TMS administration for clinical and investigational purposes.

## Introduction

Transcranial magnetic stimulation, a method generating focused magnetic field pulses over the scalp that penetrate the head, is increasingly used to non-invasively investigate brain function, and treat or diagnose conditions affecting the human brain (Rossini & Rossi, 2007). However, many details regarding the exact effect of TMS induced currents on the underlying brain tissue are poorly understood, hampering efficient and reliable application of TMS and preventing proper dose control. Hence, TMS treatment attempts and TMS based brain research can be unreliable.

This gap in our understanding of TMS has spurred several groups to attempt to model the electrical fields induced in the brain by the rapidly changing magnetic field induced by a TMS coil. To this end, realistic models of magnetic fields generated by a TMS coil were developed and empirically validated (Petrov, Mandija, Sommer, Van den Berg, & Neggers, 2017; Salinas, Lancaster, & Fox, 2007). Furthermore, the effect that the electromagnetic properties of the different tissues in the head have on the incident field produced by the TMS coil have now been modeled with finite element modeling (FEM) or boundary element modeling (BEM), thus taking into account the conductive effects of tissues in the head(Opitz, Windhoff, Heidemann, Turner, & Thielscher, 2011; Salinas, Lancaster, & Fox, 2009; Thielscher, Opitz, & Windhoff, 2011), for an overview see our recent review paper (Neggers, Petrov, Mandija, Sommer, & van den Berg, 2015). The total electric field injected by TMS inside an actual human head and brain can hence be assumed to be known to a certain extent.

How this E-field and ensuing electrical currents in the brain then evoke neuronal activation in terms of neuronal depolarization and action potentials is still a matter of considerable debate.

It is widely accepted that electric potentials oriented along the main axis of a neuron (from the soma and along the axon), results in lowest thresholds for depolarization and action potential generation (Day et al., 1989). This has led to the formulation of the macroscopic cortical column cosine (C3) model of neuronal activation by an induced E-field (Fox et al., 2004). The neuronal column is a fundamental principle of neuronal organization within the cortical sheet where the main axes of neurons are oriented in small functional columns perpendicular to the cortical sheet (Hubel & Wiesel, 1979). Hence, it is assumed that E-fields perpendicular to the cortical sheet (wherever they are deployed) are maximally effective in inducing neuronal activation. This is described by activation being proportional to the cosine of the E-field and the normal to the cortical surface at a given location (i.e. with maximally induced activation for E-fields perpendicular to a local patch of cortical sheet). Therefore, activation is based on cortical orientation relative to the induced field and not whether the local cortical surface is in a sulcus or a gyrus. Gyral crowns are usually oriented in parallel to the scalp and due to the common coil orientation also in parallel to the majority of the applied E field. This dominant orientation of the gyral crowns with respect to the applied E-fieldmake the gyral crowns less susceptible to the TMS E-field. Deeper within a sulcus, the cortical surface is oriented perpendicular to the scalp and the C3 model predicts more activation here (this effect is of course to be regarded on top of the strong reduction of induced field strength further away from the TMS coil, which in turn favours gyral crowns). Some reports claim empirical evidence for such an activation preference for the medial surface deeper in a sulcus with PET and suprathreshold TMS (Fox et al., 2004) or by the ability to explain angular dependency of coil orientation with respect to gyral orientation(Brasil-Neto et al., 1992; Kammer, Vorwerg, & Herrnberger, 2007; Petrov et al., 2017). However, the cosine model is also criticized (Bungert, Antunes, Espenhahn, & Thielscher, 2017). A few studies observed that in fact, using subthreshold concurrent TMS/fMRI, the gyral crowns are activated rather than deeper sulci (Takano 2004, Siebner 2001). Some of the most detailed modeling studies of motor cortex activation (Salvador, Silva, Basser, & Miranda, 2011) also do not show strong preference for sulcal wall activation.

Furthermore, one can argue that the cosine model, despite correctly posing that E-fields oriented along a cortical column of neurons most likely activates those neurons most effectively, ignores the fact that many of the main glutamatergic excitatory neurons in the cortical columns are interconnected horizontally by GABAergic inhibitory interneurons (Tremblay, Lee, & Rudy, 2016). One could speculate that an induced E-field polarizing and hence inhibiting such inhibitory interneurons could contribute to a maximal release of inhibition on and subsequent activation of then disinhibited cortical column neurons. This in turn would predict the opposite of the C3 model, namely activation preferentially in gyral crowns where E-fields are parallel to the main orientation of interneurons, and much less deeper in the sulci. We refer to this as the C3b hypothetical activation model.

Finally, it is possible that, due to the complex interplay between cortical columns and horizontally oriented interneurons, there is not really a clear preferential influence of the E-field orientation on the neuronal activation in a patch of the cortical sheet, which we refer to here as the ‘CE’ model.

In the present study, we put these 3 possible models of how the orientation of TMS-evoked local E-fields evoke neuronal activation in the cortical sheet local to the center of the TMS pulse to the test. We use concurrent TMS-fMRI on healthy volunteers and a pipeline of computational modeling of coil E-fields and induced fields in the brain. First, we describe the MRI image processing and volumetric parcellation to construct proper FEM models of induced currents by TMS, and detail the solving of the FEM equations. Then we proceed to elaborate a formal description and implementation of the 3 possible models for locally induced neuronal activation as introduced above. Finally, we describe the empirical concurrent TMS-fMRI technique we developed for the purpose of this study, the data fMRI analysis, and how we appraise the overlap between empirically observed with computationally derived activation.

## Methods

This study contains 2 major sections:

a. constructing a model of a TMS coil and incident evoked E-field, the ensuing induced E-field as approximated by finite element modeling taking into account 5 tissue types in the head, and several crude macroscopic models of brain activation by local directed E-fields.
b. the empirical validation of the modeled brain activation from previous section with a dedicated concurrent TMS/fMRI setup in 6 subjects

The first part of this methods section describes the modeling part in detail, whereas the 2^nd^ part describes the experimental and data analysis approach.

### Models of TMS evoked brain activation

Here we briefly describe how we model a TMS coil and how it evokes an incident E-field, which is described in more detail in a previous paper from our group. Next, we detail how we segment a T1 weighted MRI scan into 5 tissue types, and construct a volumetric mesh from those segments. After that, a description of the finite element model and induced E-field is provided. Finally, we establish 3 macroscopic models of how an E-field might interact with cortical neuronal tissue to evoke activation.

### Coil model and incident E-field

For the current paper, we adopt a model of a figure-of-8 coil with concentric wiring, and a ‘coil depth’ of 0 (meaning the coil wirings are within a plane of thickness 0). Spatial parameters: inner diameter 26 mm, outter diameter 44 mm, turn-to-turn distance: 2 mm, turns per coil: 9. For more details and the modeling followed see our previous paper by Petrov and colleages (2017), where this coil model was carefully validated using MRI phase mapping and the modeled E-field was in good agreement with the empirically measured magnetic field. See figure 1 for an impression of this model and the ensuing incident E-field (field in a homogeneous medium). The incident E-field was computed using integration of piece-wise Biot-Savart law of magnetic field generation around an electrical wire, taking into account the spiral architecture of the two coils constituting a figure of 8 TMS coil. Details are provided in Petrov et al. (2017). Dimensions of the coil wiring were derived from the actual coil used in the experiment (see experimental setup below) based on an X-ray of the coil.

**Figure 1:**
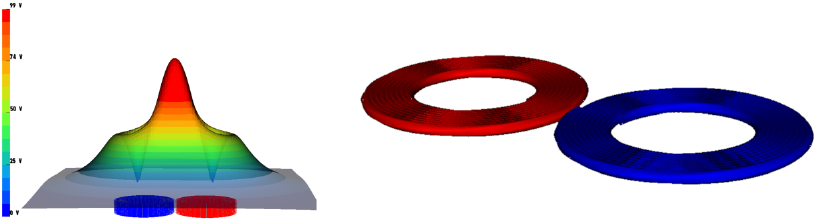
The coil model adopted, with spiral windings within a flat plane. Spatial parameters: inner diameter 26 mm, outter diameter 44 mm, turn-to-turn distance: 2 mm, turns per coil:9. For more details and the modeling followed see our previous paper by Petrov and colleages (2017).

### Deriving volumetric meshes for different tissue types from a T1-weighted MRI scan

To compute alterations of the total E-field caused by inhomogeneity of the tissues in the head and brain, we adopt finite element modeling of an E-field in an in homogeneous medium (for details, see Neggers et al 2015). For FEM to work properly, it is paramount that the spatial layout of underlying tissues such as skin, skull, cerebral spinal fluid (CSF) and gray and white matter (GM, WM) are described volumetrically in such a way that they capture the electromagnetic properties (especially electrical conductivity) well.

In order to do this, we segmented a T1 weighted MRI scan (TR/TE of 10.0/4.6 ms, a flip angle of 8°, voxel size of 0.75×0.75×0.8 mm3, scan duration of 11.3 min, 225 slices with a slice gap of 0 mm) using the ‘unified segmentation’ algorithm as implemented in SPM12 with Bayesian prior tissue maps for the 5 aforementioned tissue types and air (Ashburner & Friston, 2005). This generally results in viable segmentation for each major tissue type in the form of a probability map where voxel values indicate the probability of belonging to a certain tissue class. For each voxel in the volume, it was then determined what the most likely tissue type was (i.e. which tissue probability map had the highest value at that location, where the ‘winning’ tissue type had to be at least 0.3 likelihood). This tissue type was stored in a new volume as an index. The CSF index values were stored in first and dilated by a voxel, such that no holes were present between GM and CSF, then in the same fashion the WM values. The other tissue indexes where added after that. The indexes where: 1 = GM, 2 = WM, 3=CSF, 4 = Skull/Skin, 5=air. Note that skull and skin were treated as one tissue type, as mainly the inner boundary of the skull is relevant for current flow in FEM.

We inspected the indexed image visually slice by slice for every single participant included in the study, and occasionally manually removed voxels erroneously classified as GM deeper inside sulci that ‘clogged’ a sulcus (causing the GM voxels across a sulcus to ‘connect’). Such ‘clogging’ would have caused the 2 banks of a sulcus to artificially connect in a computed volumetric mesh. See figure 2a for a visualization of this manual editing process.

**Figure 2a.**
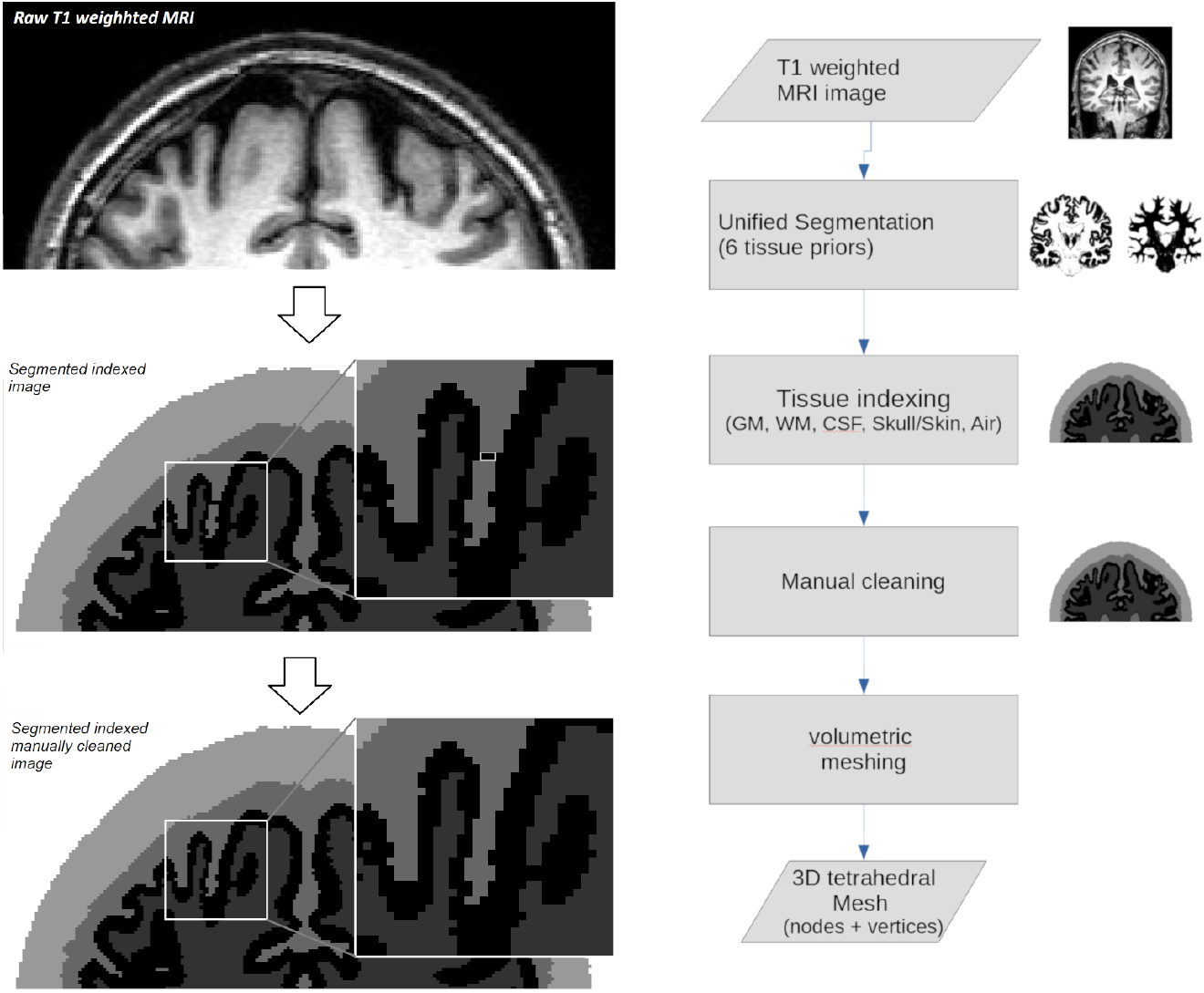
(Left) Steps in tissue segmentation from unprocessed T1 weighted MRI to clean tissue maps for gray matter colored black), white matter (dark gray), cerebral spinal fluid (medium gray) and merged skull/skin (light gray). Figure 2b (Right) Overview of all major steps in our 3D meshing pipeline. First step is to acquire T1-weighted MR image. Second, we need to segment (split) the image in discreet/binary areas per tissue type. Step three will involve any manual cleaning of the segmentations before attempting building a 3D tetrahedral mesh. Initially the 3D mesh will comform ideally to the voxel mesh of the original image. Only after some iterations the mesh will be optimized to better fit the border areas, where two or more tissues stand adjoin. In the process, the geometry is guaranteed to maintain water tight properties, that is all isolated volumes outer surfaces will form a closed surface with no holes.

The binary tissue index maps were first isolated per compartment and pre-processed (explained below) before fetched as input for the 3D meshing algorithm.

3D meshing is the process of describing a volume (in this case tissue type) with elementary shapes, in our case tetrahedrons. To guarantee rather smooth boundaries on the interface between different compartments, we first adopted an iterative smoothing routine of alternating inflation-deflation operations in image space. Those images processed by our meshing algorithm one per tissue type to produce the final FEM mesh using Cleaver2 : A Multi Material Tetrahedral Meshing Library and Application (Scientific Computing and Imaging Institute (SCI), University of Utah, The USA, https://github.com/SCIInstitute/Cleaver2). For more details on this procedure, see Petrov et al (2023).

In figure 2b the entire MRI image processing pipeline leading to a 3D meshed head is depicted and further elaborated upon.

### Finite Element Model of TMS evoked field in the head

The FEM model computing the induced field in the head generally is posed as in formula [1]:

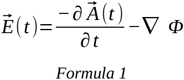

There are two components contributing to the total electrical field induced by TMS, 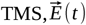. In [1], 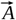 is the vector potential of the magnetic field 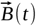 directly generated by the TMS coil, ☞ the scalar potential generated in the tissue, the gradient of which gives the secondary component to the total electrci field Et. The first term on the right hand side of [1] reflects the component of the E field directly generated by the fact that a changing magnetic field flux is applied over a volume, and would also be present in vacuum. The scalar potential ☞ is more complex, and arises from charge accumulations in the brain that will be created as currents start to flow. Currents will experience resistance when flowing through a complex conducting medium. When the magnetic field 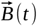 induced by the TMS coil and the conductivities of the tissues in the head are known, the gradient of the scalar potential can be solved using the finite element method (FEM) of the equations governing accumulating charges in the head *per mesh element*.

We used SCIrun v4.7 with our own custom modules (see Petrov et al 2023 for details including open sources of our own models) to solve the FEM equations and derive the scalar potential ☞.

### Macroscopic models of brain activation in the cortical sheet

With a total E-field determined as described above, it is still not known how neurons in the cortical sheet will respond to such a field and invoke brain activation. Whereas it is at present extremely difficult to describe this process in detail at a microscopic single neuronal level, there are sensible assumptions one can make at that level (Fox et al., 2004). We looked at three ‘metrics’ as possible cortical activation predictors. Each ‘metric’ represents a macroscopic model of how the total E-field experienced by a small patch of the cortical sheet acts on pyramid cells within that patch, reflecting knowledge of the microscopic structure of the cortical sheet. Per metric, we also developed a method to implement such metrics on a surface mesh of the cortex, allowing us to experimentally validate these models with concurrent BOLD-fMRI on the same subjects for which the cortical mesh, FEM and the three metrics were determined. Figure 3 illustrates each of the metrics of neuronal activation by a local induced E-field: C3 (activation induced at descending tracts), C3b (activation induced by interpyramidal connections), CE (strength of total E-field). Also, the approximation of each activation metric by a triangular surface mesh has been depicted for each metric at several locations on a gyrus.

**Figure 3.**
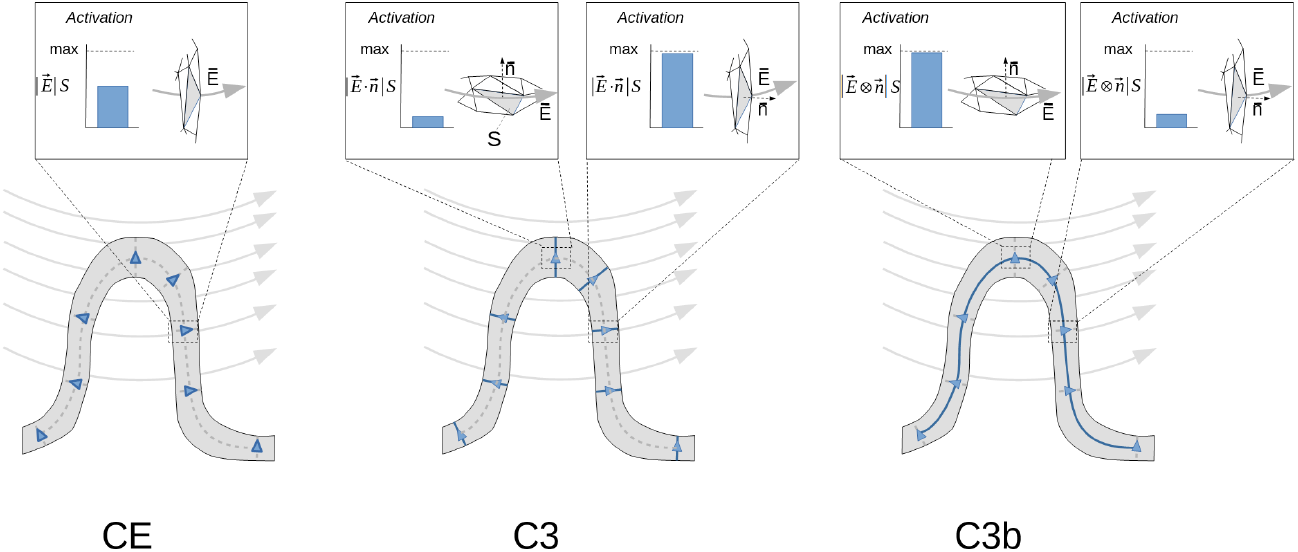
The 3 alternative models (or metrics) we adopt for how induced E-field interacts with the cortical sheet are depicted here, with on top the expected activation magnitude as a function of angle between local E-field and surface normal. From left to right: C3 (maximal activation for local E-field orthogonal to the cortical surface), C3b (maximal activation for E-fields parallel to a local surface), and CE (simply the magnitude of the local E-field independent of angle with respect to the surface). An example of activation at the crown of the gyrus and at a location deeper within the sulcus is provided to highlight the characteristics of each metric.

#### Metric C3

It has been established that brain activation, as estimated by proxy through measured electromyograms (EMG) as a response to TMS on the motor cortex (motor evoked potential or MEP), is affected by the direction of the underlying central sulcus (Basil Neto et. al., 1992). Currents induced perpendicular to the central sulcus evoke maximal MEPs, but to evoke MEPs with the same amplitude, a larger current magnitude was needed for currents evoked parallel to the central sulcus. Similar observations were made for the direction of current with respect to major sulci in the visual cortex in terms of evoked phosphene thresholds (a visual illusion). Such observations led to assumptions that neuronal axons oriented perpendicular to the cortical sheet, potentially descending tracts, cause the majority of evoked activations by TMS, and hence are stimulated optimally with currents perpendicularly to the cortical sheet. This has been dubbed the ‘C3’ metric for the relationship between currents and neuronal activation(Fox et al., 2004). See figure 3 for a graphical depiction.

When approximating the cortical surface with a triangular surface mesh, as depicted in figure 3 per metric, a surface patch with a set of triangles would experience the total ‘activation’ of the inner product between the surface normal of each triangle and the total E-field through that triangle (maximal for a local E-filed aligned with the surface normal), multiplied by the surface of that triangle S, yielding the term 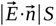, summed over all triangles in a patch. The absolute value is used because we assume a symmetric interaction for inward and outward E-fields (see below), due to the bi-phasic stimulator we employ

However, there is ample debate about this metric (Thielscher et al., 2011), and we hence also adopted other potential metrics that might explain neuronal activation induced by a local current.

#### Metric C3b

It could also be that local horizontal connections between pyramid cells in the cortex, through inhibitory interneurons (Tremblay et al., 2016), produce most of the activation induced by local fields. In such a case, currents in parallel to a cortical tissue would evoke maximal neuronal activation (or suppress it maximally). This metric we dub C3b is also sketched in figure 3 below.

For C3b, a cortical sheet approximated by surface with a set of triangles would experience the total ‘activation’ of the length of the outer product between the surface normal of each triangle and the total E-field through that triangle (maximal for a field perpendicular to the normal), again multiplied by the surface of that triangle S, yielding the term 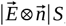, summed over all triangles in a patch. The absolute value is used because we assume a symmetric interaction for inward and outward E-fields (see below).

#### Metric CE

Finally, we adopt the notion that perhaps the mix of neuron types found in the cortex is too complex for such a simple and crude neuronal metric of activation, and that the best predictor for activation is simply the magnitude of the local E-field without taking into account its direction. We describe this metric as ‘CE’. A surface patch representing the cortical sheet would for this simple metric contribute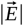 *S*per triangle, where S is the surface of each triangle.

Note that one could argue that the E-field’s polarity also should be taken into account. For example, for metric C3, this would mean that currents pointing inward from CSF into a sulcus, would evoke a negative (or minimal) signal, and pointing outwards a maximal signal, which would equate to 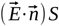. The same would then apply to C3b metric. However, while this idea could hold for a monophasic TMS pulse shape, we validate these models with a TMS device evoking biphasic pulse shapes, there is a current in opposing directions in short succession per stimulus. Therefore we can not distinguish between a symmetric or assymetric coupling between E-field and neuronal sheet in our validation experiments, and hence we assumed a symmetric interaction and implement this notion with the absolute value of the inner product between the E-field and the surface normal, or 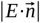 for C3, and in a similar fashion for C3b.

In figure 3, the C3, C3b and CE metrics are depicted, including how to compute them for a triangulated surface mesh we use to describe the cortex with for sake of simplicity and to compare it with fMRI BOLD data from our validation experiment.

In figure 4 it is depicted how our implementation of these metrics lead to surface represented predictions of neuronal activation induced by TMS for an actual surface mesh of the cortical sheet of one of our participants.

**Figure 4.**
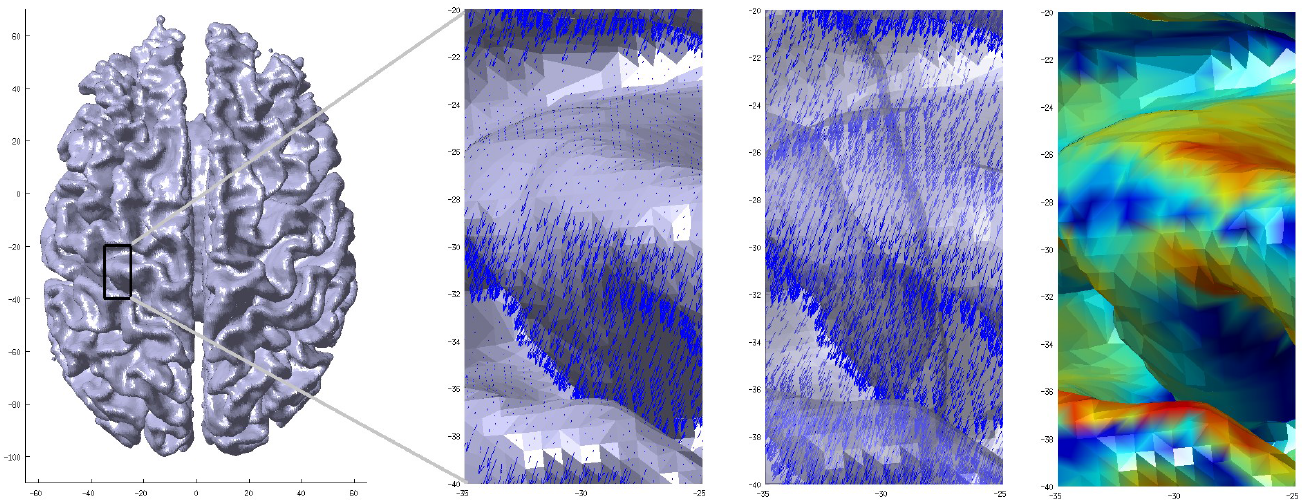
An illustration of how the C3 metric is derived. From left to right: cortical surface of one of the participants, rendered halfway the pial surface of the cortex and the gray-white matter boundary; zoomed in view of the E-field vectors induced by TMS for each triangular surface patch (inward pointing vectors not shown); same figure as to the left, but now with transparent surface to allow view of inward pointing E-field vectors; same surface patch color coded with C3 activation metric computed from normal of each triangular surface patch and corresponding E-field vector (red=high and blue=low (0)).

### Validation of modeled brain activity patterns with concurrent TMS/fMRI

In 6 healthy individuals, we evoked brain activation with TMS inside a 3T MRI scanner. To this end, a dedicated concurrent TMS/fMRI setup was used. The human primary moto-cortex area-M1 was targeted with single pulse TMS at 2 intensities, and ensuing fMRI time series acquired. It was investigated whether evoked BOLD activation near the stimulation target matches modeled ‘activity’, by using FEM-derived total E-fields based on individual brain morphology as described in the previous section, and the 3 aforementioned activation ‘metrics’, confined in the gray matter. It is analyzed which of the metrics results in the best match with measured fMRI BOLD responses to TMS.

Details of the experiment and comparative analysis are provided below.

### Participants

6 healthy individuals (1 m, 5 f, mean age 22.8 yr) were included in this study. All participants were screened for contraindications against MRI or TMS, and provided written informed consent. Procedures were approved by the Medical Ethitcal Committee (METC) of the UMC Utrecht (protocol number 16-469).

### Concurrent TMS/fMRI experimental setup

This section provides a summary of the dedicated concurrent TMS/fMRI setup used in this study. Details can be found in 2 of our previous studies where the same setup was used(de Weijer et al., 2014; Vink et al., 2018).

All MR sequences were performed in a 3T MR scanner (Achieva, Philips Healthcare, Best, The Netherlands).

### Experimental design

The experiment consisted of a short intake session ((f)MRI-only) and a TMS session (concurrent TMS-MRI) on a different day.

During the intake session, a normal 8 channel SENSE enabled head coil was used and subjects were placed in a standard position during head scanning. A 3D T1 weighted anatomical scan was acquired with a TR/TE of 10.0/4.6 ms, a flip angle of 8°, voxel size of 0.75×0.75×0.8 mm3, scan duration of 677 s, 225 slices with a slice gap of 0 mm. Next, a single-shot echo-planar imaging (EPI) scan was acquired with 250 dynamics, a TR/TE of 2,000.0/23.0 ms, flip angle of 70°, voxel size of 4×4×4 mm3, a scan duration of 510 s and 30 slices with a slice thickness of 3.6 mm and a slice gap of 0.4 mm. Participants moved the thumb of the right hand after hearing of an auditory cue. The functional data was analyzed to determine the representation of the thumb area in the left M1 as determined with BOLD fMRI. Preprocessing and statistical analysis are described in the data analysis section below. Also during the intake session, the resting motor threshold (RMT) was determined by stimulating the primary motor cortex while increasing the TMS stimulator output until a response in the APB muscle was visible in 5 out of 10 TMS pulses[26].

The TMS session started with neuronavigation to determine the coil position overlying the individual participants M1, using The Neural Navigator by Brain Science Tools BV, the Netherlands (www.brainsciencetools.com). Neuronavigation was performed outside the MRI scanner in a separate room, and M1 was marked on a bathing cap the participants were wearing. The T1 weighted anatomical scan from the intake session was used for neuronavigation. Based on the 3D brain surface, the M1 target was obtained from the activation map acquired during the intake session, and used as a target during neuronavigation. Thereafter, the participant was placed in the MRI scanner with the concurrent TMS/MRI setup and underwent a combined TMS-fMRI sequence in which single TMS pulses were delivered to M1. Finally, a T2-weighted scan was acquired to retrospectively verify TMS coil placement. For details on coil position reconstruction from the T2 weighted MRI, see Vink et al (2018) and De Weijer et al (2013).

After successful TMS coil positioning, two sequences were acquired. First, a T2-weighted scan with a TR/TE of 13,609.0/80.0 ms, flip angle of 90°, voxel size of 2×2×2 mm3, scan duration of 218 s. This was done by attaching 6 custom made markers (small capsules filled with water) to the back of the TMS coil (Fig. 2B), which appear hyper intense on the T2-weighted scan (Fig. 1). Second, a single-shot EPI sequence was acquired with 500 dynamics, a TR/TE of 2,000.0/23.0 ms, flip angle of 70°, FOV of 256×119.6×208 mm3, matrix of 64×63, voxel size of 4×4×4 mm3, scan duration of 1020 s, 30 slices with a slice thickness of 3.6 mm and a slice gap of 0.4 mm. During the EPI sequence, single pulses of TMS with an intensity of 115% RMT were interleaved with pulses with an intensity of 60% RMT. TMS pulses were delivered with a random interval of 5 to 8 dynamics (10 to 16s) to avoid habituation. It has been shown that the MRI static magnetic field affects the flow of current through the TMS coil, which reduces the TMS magnetic field amplitude[28]. Therefore, we decided to set the RMT at 115% instead of 110% for suprathreshold stimulation.

### fMRI data analysis

fMRI time series data were analyzed using SPM12 and several custom matlab scripts, all running in a Matlab R2014a environment (Mathworks Inc., USA).

First, all EPI volumes were inspected to determine image quality and to identify the presence of potential artifacts. This revealed small random deflections from the baseline signal level in a single slice of a few functional volumes per time series acquired during the TMS session. A small number of artifacts were present in most of the time series data, most likely caused by manually changing the TMS device intensity in between TMS pulses (and hence charging the stimulator capacitors). These deflections are short (one sample) and can only be observed in the vicinity of the TMS coil. All slices of the realigned EPI scans were automatically scanned for the presence of a sharp peak in the average grey matter signal with a custom algorithm to detect distortions. The distorted slices were then interpolated based on the BOLD signal in the previous and next slice with custom Matlab code. An average of 71 slices were interpolated per participant, out of a total 30 slices and 500 volumes.

Likely, removal of the large artifacts due to coil charging does not rule out smaller fluctuations that might still be present in the data (see results section).

Next, the fMRI and anatomical MRI data were preprocessed. The functional MRI time series were corrected for head movement using least-square minimization and rigid body transformations. Next, the time series data was coregistered (not resliced) to the T1 weighted anatomical scan using a rigid body transformation and a mutual information registration method from SPM12. The images were then smoothed with 8mm full-width at half maximum.

Next, a GLM approach was adopted to compute the amount of activation observed on average for the 2 intensities of TMS stimulation. The generalized linear model (GLM) included two events: single pulses of 115% RMT and 60% RMT. The BOLD response was modeled with the canonical hemodynamic response function (HRF) and its first-order derivative. Two nuisance regressors were included in the analysis: the average BOLD signal in the white matter and the CSF. BOLD signals were filtered with a high pass filter of 80Hz before construction of the GLM. Statistical images were constructed based on the contrast between TMS pulses of 115% RMT and TMS pulses of 60% RMT using an T-statistic with a threshold at P < 0.05, family wise error (FWE) corrected (Penny et al, 2011)

Finally, the activation maps (T-maps and contrasts) were resliced to the lattice of the T1-weighted anatomical scan for proper comparison with the model values, also expressed in this lattice (see below).

Note that the above analysis does not include a normalization to MNI space, the data stays represented in so called ‘native space’ as they were oriented in the MRI scanner. This was needed as the coil, FEM and activation metric modeling described above had to be performed in native space as well, and it is our goal to compare BOLD activation with models. As the participants had to maintain a tilted head angle with respect to the normal posture to allow placement of the TMS coil (see ‘experimental setup’ above), this native space data is less suitable to display activation maps graphically as slicing would be under an oblique angle. The activation maps and anatomical scan were normalized to MNI space using the normalization parameters from the unified segmentation step described in the session on volumetric meshes above. This was purely done for visualization purposes of the fMRI BOLD data, all analysis are performed in native space.

### Comparison of fMRI activation maps with modeled brain surface ‘activation’

fMRI activation maps, analyzed as the GLM regression coefficients (contrast maps) reflecting relative signal change induced by TMS as described above (high intensity versus low intensity), need to be compared to the surface triangulation based modeled ‘activation’ maps described in the previous section. The fMRI maps are expressed in a rectangular grid compliant with the Nifti v1.1 data specifications (according to the T1-weighted anatomical scan because of the reslicing, see previous section), but the modeled ‘activation’ comes as a value per triangle on the extracted brain surface. Hence, we need to convert the modeled activation from a triangular surface representation to a matching rectangular grid. We accomplished this as follows. While iterating over all triangular faces representing the brain surface and having an associated model ‘activation’, we computed for each triangle which grid points in the lattice of the T1 weighted anatomical scan are closer than 1.5mm to the triangle (shortest orthogonal distance to the triangle). Those lattice points (voxels) were given the vale of the model ‘activation’ associated with that triangle as explained in the previous section. All voxels not in the vicinity of any triangle are given the value zero.

This way, we obtain an ‘activation’ map in the same image lattice as the analyzed fMRI bold activation maps so they can be compared. In this created image, all activation values are constrained in the gray matter by virtue of the above procedure. An example of such a model activation map is given in figure 5 below.

**Figure 5.**
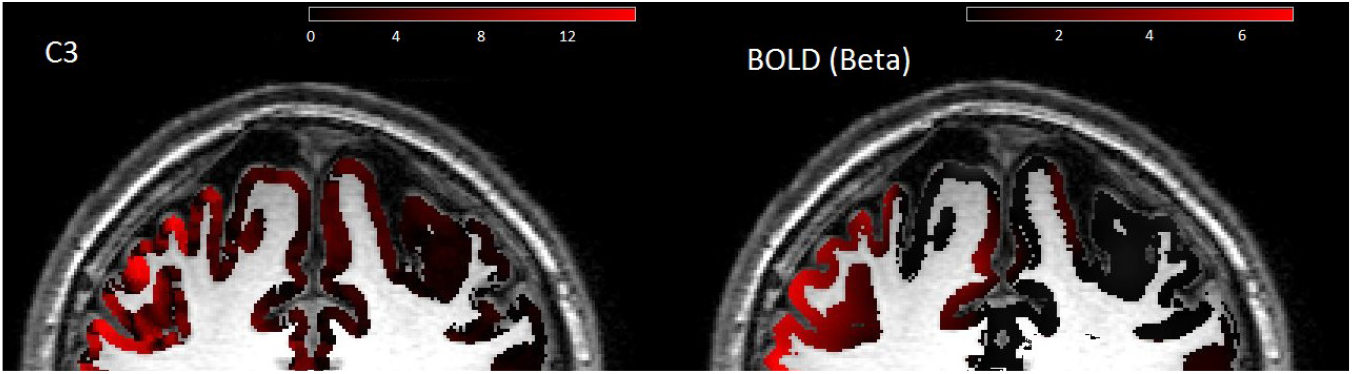
After the activation metric C3 is computed for the cortical surface (actually a layer halfway in the gray matter), a 3mm thick layer of voxels is ‘tagged’ around each triangular surface patch (see Figure 4) with the value of that nearby ‘activation’. This activation ‘map’ in voxel space allows for direct comparison with measured BOLD fMRI activation maps, shown in the right panel for the same subject and same stimulation location.

For a proper comparison, the fMRI activation maps, represented by the contrast between high and low intensity stimulation (difference between *β*_high_ and *β*_low_), are also masked by the gray matter indexed map obtained from tissue segmentation (see section on ‘*Models of TMS evoked brain activation’*) such that voxels outside the gray matter are set to 0. This procedure assures the modeled ‘activation’ can be compared to the measured BOLD activation in a rectangular lattice (a ‘voxel based’ comparison).

The overlap between the measured BOLD activation map and the modeled activation (for each metric) forced into a rectangular lattice representation as described above is assessed as follows. First, both the BOLD fMRI activation map and the modeled ‘activation’ map are thresholded at a ratio of 0.3 (30%)of the maximum activation value, where values below the threshold are set to 0 and above threshold to 1. Then, the Dice Sörenssen coefficient (DSC) is used to computed overlap between the two images. This DSC index originally stems from agriculture, and was used to compare vegitation on fields, and has later been adopted for comparing binarized medical tissue images in segmented images, including brain segmentations (Baselice, Ferraioli, & Pascazio, 2015; Zou et al., 2004). This index is straightforward, and computes the fraction of overlapping voxels (3D image pixels) between 2 binarized images X and Y in relation to the total volume of the segmented image, as follows:

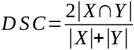

This index yields a maximum of 1 when the two images compared are identical, and 0 when they are completely non-overlapping.

DSC’s are computed for the following 3 comparisons:

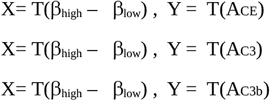

where *β*_high_ – *β*_low_ is the contrast between high and low TMS induced BOLD activation, ACE the model activation for the CE metric (total field) as explained in the section on modeling above. T is a thresholding function where:

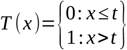

The threshold value t equals 0.3 x the maximum value of the map that is thresholded.

The DSC is computed for voxels within a spherical region of interest (ROI) around the location of maximum activation in the CE model. This is done to exclude voxels in regions further away from the motor cortex, that might be activated through connections, or auditory signals due to the fact that the supra-threshold TMS pulse still evokes a louder clicking sound than the sub-treshold TMS pulse. This way, the most direct comparison possible is made between directly activated cortical layers and modeled activation in cortical layers.

## Results

### TMS induced BOLD activation

For 6 participants, the TMS BOLD activation was analyzed as described in the methods section. The regression coefficients for high and low TMS machine output were contrasted, and overlayed on the anatomical MRI of each participant. In figure 6, the fMRI activation for high versus low machine output TMS is presented in a slice through the motor cortex for 2 participants.

**Figure 6.**
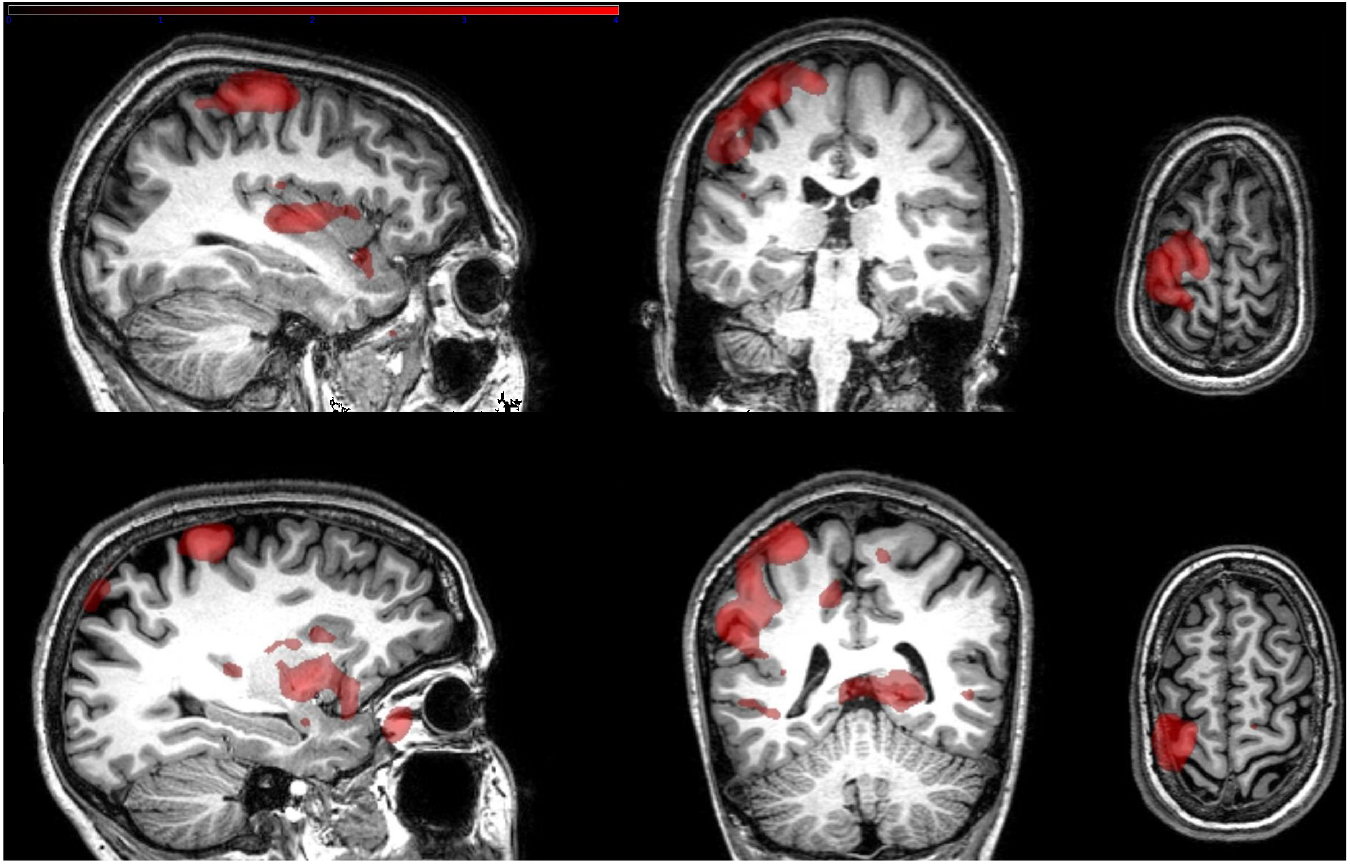
TMS evoked fMRI BOLD activation maps for two participants, following single pulse TMS stimulation on left M1 inside a 3T MRI scanner. Contrast: supra vs subthreshold TMS. Clear activation maps are observed near the stimulated M1.

There is a focus of activation near the targeted region for most participants. Targeting was not always successfully maintained while the setup was in the MRI bore, sometimes regions near but not exactly on the motor cortex were targeted with the isocenter of the TMS incident field (also see figure 7 with 3D renderings). Activation in the targeted motor cortex area tends to be in the more superficial parts of the cortical sheet (the gyral crowns) and not so much in the deeper parts of the sulci.

**Figure 7a.**
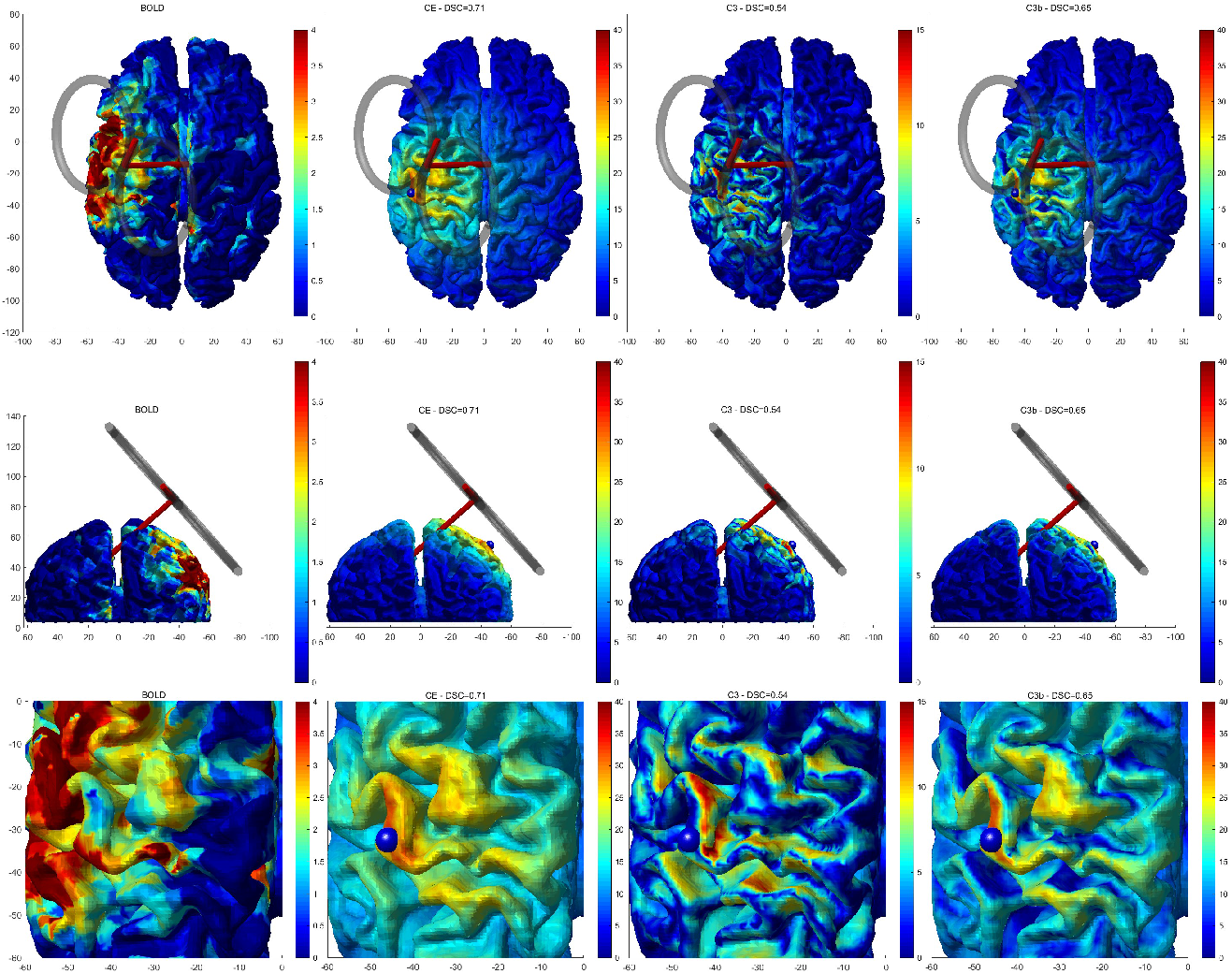
For one of the particpants, the TMS evoked BOLD activation is shown (left panels), alongside the modeled ‘activation’ based on FEM modeling of the evoked current in this particular indivudual 3D meshed brain, and the three alternative ‘metrics’ for interaction of evoked E-field with the cortical tissue (from left (2nd column) to right: CE, C3 and C3b. Note that for this participant, a TMS artifact was still visible near the left temporal lobe, most likely due to recharing the condensator of the stimulator, that affected the image somewhat for some participants. We therefor only evaluated an area proximal to the stimulation site. The blue sphere denotes the intersection of the main figure of 8 central axis pointing down from the coil surface, representing the location of maximum evoked E-field (shown as a red cyclinder in the two top rows). A simplified model of the TMS coil (2 rings) is also rendered in the top 2 rows to illustrate coil position and location in the scanner (which was established with a T2 scan after fMRI acquisition, see methods section). The bottom row shows a zoomed in view near the stimulation site. BOLD activation and the resulting ‘metric’ for modeled activation is color coded for values at the cortical surface (colorbars represent modeled ‘activation’ (arbitrary units) or T-values for the fMRI BOLD maps in the left column).Also, the overlap coefficients (DSC or Dice-Sørensen Coefficients) between model maps and observed fMRI BOLD maps are given in the titel of each panel.

**Figure 7b.**
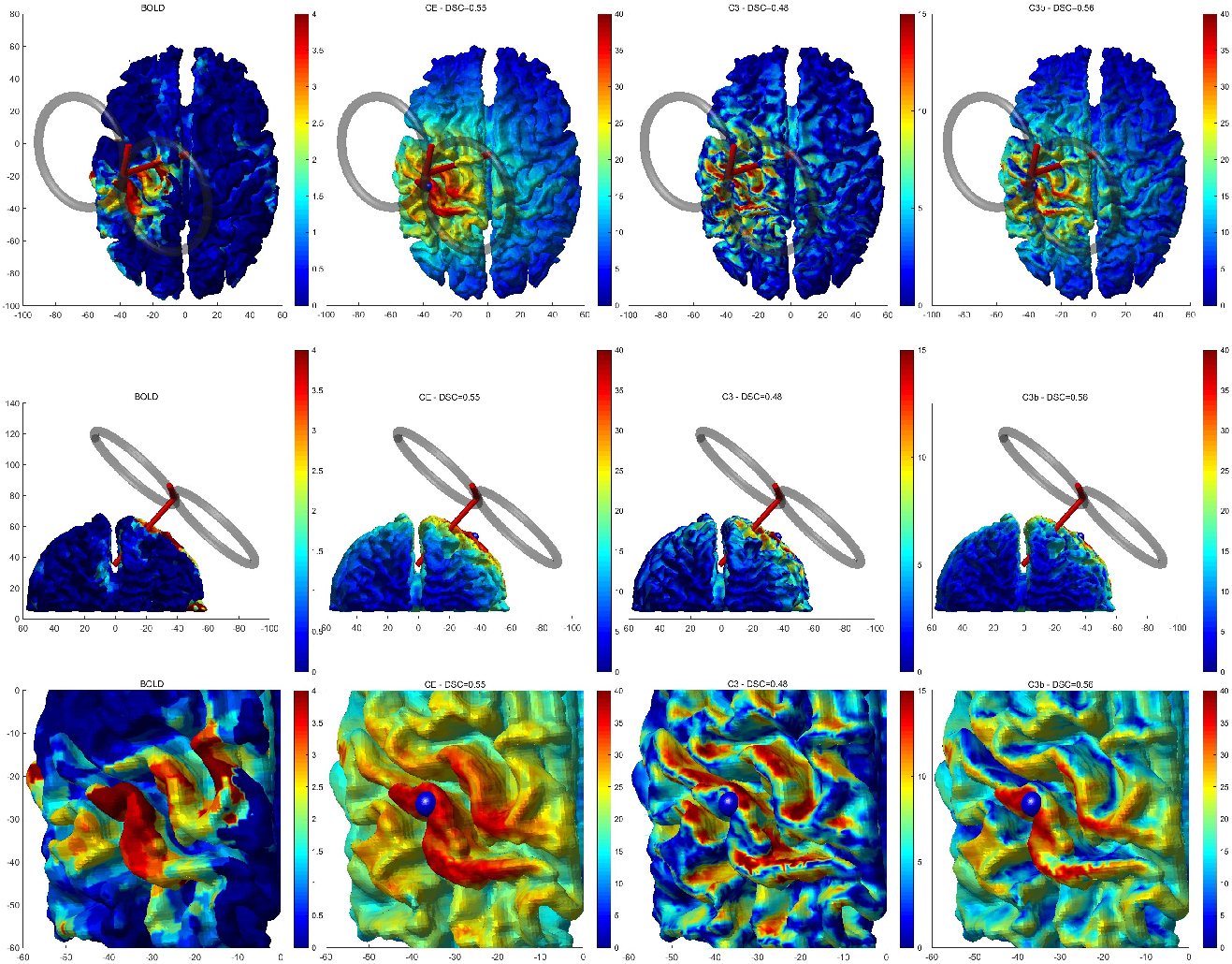
The same information as in figure 7a but for another participant.

Also, in the left temporal lobe, there is a larger cluster of activation for some participants, that extends into non-neuronal tissue outside the brain. We believe this is an artifact due to the manual alteration of intensity during the experimental session, as explained in the methods section (‘fMRI activation analysis’). In the 3D renderings of BOLD activation below, it can be seen that these potential artifacts are close to the coil wing near the temporal lobe, especially when the coil deviated from its ideal position parallel to the scalp. For this reason, the BOLD activation was only evaluated within a ROI of 4cm around the maximum model activation (generally near the maximal field center line of the coil, see methods section), largely excluding this artifact from the assessment.

### Comparison of model and BOLD activation patterns

As described in the methods section, the DSC was computed to compare overlap of model and BOLD activation for each of the 3 metrics. See figure 7a and 7b below for 2 participants, where a cortical rendering is presented of TMS evoked BOLD activation and modeled activation for the 3 metrics. In the title, the DSC overlap coefficient between measured BOLD and modeled activation is given for each metric. The cortical renderings for all other participants are given in the supplementary material in a similar fashion.

As it can be seen from figure 7, for both subjects there is some spatial resemblance between the BOLD activation and especially the activation following the ‘total induced field’ metric CE and the ‘parallel’ activation metric C3b model, that all show activation primarily in the gyral crown. The C3 metric results in ‘activation’ that has maximal values somewhat deeper inside the sulci. This pattern of resemblance is reflected in a lower DSC for the comparison of activation following the C3 metric with respect to the BOLD activation.

In table 1 below, the DSC coefficients for all 6 participants is given for all 3 metrics. There is a trend for a lower DSC for overlap between measured BOLD activation and modeled activation due to C3 as compared to the other 2 metrics (T=1.722, p=0.058).

**Table 1.**

| *DSC* | *CE* | *C3* | *C3b* |
| --- | --- | --- | --- |
| S1 | 0.7108 | 0.5392 | 0.6487 |
| S2 | 0.5453 | 0.4757 | 0.5634 |
| S3 | 0.4685 | 0.4416 | 0.3245 |
| S4 | 0.4849 | 0.4594 | 0.4303 |
| S5 | 0.5849 | 0.4672 | 0.5211 |
| S6 | 0.4482 | 0.4083 | 0.4064 |

## Discussion

We managed to successfully stimulate 6 participants with TMS on or near the motor cortex while acquiring BOLD fMRI data. Observed BOLD activation tends to be primarily focused in the more superficial parts of the cortical sheet, the gyral crowns.

After segmentation and volume meshing of the anatomical T1 weighted MRI scan, calculation of the incident field of the TMS coil and estimation of a Finite Element Model, we obtained a realistic individual total induced field pattern for the same 6 participants. We applied 3 putative models of how induced E-fields induce neuronal activation following different assumptions on cell types that are sensitive to TMS induced fields: the well known C3 cosine model assuming neurons oriented along cortical columns perpendicular to the cortical field attribute the majority of the activation, an alternative model C3b where horizontally directed interneurons contribute most to the activation, or a simpler model CE where the magnitude of the total induced E-field activates neurons, irrespective of the direction of the induced E-field.

When directly comparing the observed activation with modeled activation in the cortical sheet, most overlap was found for the CE model where no directional preference was observed. A lower overlap was found for the C3b metric of neuronal activation, and the worst performing metric was the well known C3 metric based on cortical column orientation.

When inspecting the measured BOLD activation and the modeled activation patterns on the cortical surface, as depicted in figure 5, the above finding seems to be largely explained by the fact that the BOLD observation is constrained to the more superficial part of the gyral crowns, rather than deeper in the sulci. Both the CE and C3b model predict more superficial ‘activation’, and hence show most overlap with the measured BOLD activation. The CE and C3b models did not differ substantially among each other.

These findings seem to match the findings of (Bungert et al., 2017; Opitz et al., 2011, 2011) best. Their brain modeling results also exhibited strongest superficial rather than sulcal induced currents, which they attributed not only to the incident field strength decaying rapidly below the coil surface, but also the resistance caused by the narrow passages between the gyral crowns and the inner side of the dura mater that constrains the flow of cerebral spinal fluid. It has to be noted here that in the aforementioned studies, induced currents were assessed directly, and not through macroscopic neuronal activation metrics as attempted here. Although Bungert et al did not directly compare their modeling results to functional MRI activation in the same subjects as the present study attempted, they did analyze functional MRI literature on motor activation and concluded from that literature that indeed fMRI activation tends to be more superficial than deeper withing the sulci for motor related paradigms. The assessment by as somewhat hampered by the necessity to use MNI space averages and infer sulcal activation maps from such reported coordinates, which can be quite different from native space non-normalized brain space, the results point in the same direction as the present study that did compare modeling work with activation directly in 6 participants.

The present study had several important limitations that need to be discussed. First, the BOLD fMRI experiments were, for some subjects, plagued by an artifacts that seemed related to the manual alterations of TMS machine output during the experiment. We were able to remove the larger spikes but some more subtle activation that correlated with the super versus subthreshold stimulation paradigm adopted here, was still present in the activation contrast as can be seen in figure 7. This artifact mainly occurred near the temporal lobe directly below one coil wing. For the BOLD and model activation overlap assessment, we focused on activation near the motor cortex using a spherical region of interest approach in order to exclude the artifact from our results.

Second, for computational reasons we merged the skin and skull tissue segments. This greatly simplified the volumetric meshing of those tissue volumes as the boundary between them was noisy in our data, and unsuitable for FEM. We might, however, have overestimated the secondary currents as modeled by FEM (right-most expression in equation [1]) as a result.

Third, the overlap assessment of BOLD with modeled ‘activation’ by the DSC value is probably simplistic, for several reasons. The first reason is that BOLD activation evoked by TMS reflects the full pattern of neuronal activation, both local to the TMS evoked maximal field, as well as local and distal responses to such neuronal activation through connections, through secondary activations caused by TMS such as auditory or somatosensory signals (despite the fact that we contrasted high vs low intensity TMS there still is a difference between both conditions), and possibly through reafferent activation from evoked muscle activation due to suprathreshold stimulation. The modeled activation on the other hand only reflects directly activated cortical tissue, and completely disregards activation through local and distal connections. Nevertheless, one can safely assume that also the BOLD activation at least reflects the same type of direct neuronal activation as the modeled direct activation, and hence shows a larger degree of overlap for correctly modeled activation.

In conclusion, the present study observed that at the moment total TMS evoked electric field assessments as modeled by FEM are sufficient to describe neuronal activation that occurs mainly in the gyral crowns, for both BOLD fMRI and modeled activation. Activation preference for currents running in parallel with the cortical surface performed equally well, but the well known C3 metric where the orientation of cortical columns perpendicular to the gyral wall are supposed to explain the largest part of TMS evoked activation (Fox et al., 2004), performed worst of all 3 activation models tested. Further research with higher resolution imaging techniques should be able to better discriminate between total electric field and currents parallel to the cortical sheet, or more detailed cortical layer models.

## Acknowledgements

This work was supported by the ENIAC grant entitled DeNeCor ‘Devices for NeuroControl and NeuroRehabilitation’.

